# Causes and consequences of mitochondrial proteome size-variation in animals

**DOI:** 10.1101/829127

**Authors:** Viraj Muthye, Dennis Lavrov

**Affiliations:** Bioinformatics and Computational Biology Program, Iowa State University, 2437 Pammel Drive, Ames, Iowa 50011, USA; Department of Ecology, Evolution and Organismal Biology, Iowa State University, 241 Bessey Hall, Ames, Iowa 50011, USA

**Keywords:** mitochondria, proteome, mitochondrial-targeting-signals, animal

## Abstract

Despite a conserved set of core mitochondrial functions, animal mitochondrial proteomes show a large variation in size. In this study, we analyzed the putative mechanisms behind and functional significance of this variation using experimentally-verified mt-proteomes of four bilaterian animals and two non-animal outgroups. We found that, of several factors affecting mitochondrial proteome size, evolution of novel mitochondrial proteins in mammals and loss of ancestral proteins in protostomes were the main contributors. Interestingly, gain and loss of conventional mitochondrial targeting signals was not a significant factor in the proteome size evolution.

## 1 Introduction

Mitochondria, membrane-bound organelles present in most eukaryotic organisms, are involved in a number of cellular processes, including oxidative phosphorylation, Fe/S cluster biosynthesis, amino-acid and lipid metabolism, and apoptosis [1, 2]. These tasks typically require more than a thousand proteins, the vast majority of which are encoded in the nuclear genome and imported into organelle [3, 4, 5]. Thus, analysis of nuclear-encoded mitochondrial proteins (mt-proteins^1^) is essential for understanding mitochondrial function and evolution.

Several approaches have been used to estimate the composition of mitochondrial proteomes (mt-proteomes^2^), which can be roughly divided into those directly extracting proteins from mitochondria and those that infer their mitochondrial localization using a variety of analytical methods [6]. The former methods include mass spectrometry (MS) and microscopy. The latter include identification of targeting signals, homology search, co-expression analysis, and phylogenetic profiling. Since all these techniques have their pros and cons [6], integrated approaches have been advocated and applied to elucidate some mt-proteomes, as in the case of human and mouse MitoCarta [7]. Despite these advances, the number of experimentally determined mt-proteomes remains small.

Mt-proteomes have been characterized in several eukaryotic species, including plants [8, 9, 10, 11], fungi [12], animals [7, 13, 14, 15] and protists [16, 17, 18, 19]. Within animals, mt-proteomes have been experimentally determined in a few model species: mammals (human, mouse and rat [7, 20]), arthropod (*Drosophila melanogaster*) [14] and nematode (*Caenorhabditis elegans*) [13]. These studies revealed a large size difference between mammalian mt-proteomes and those of protostome animals (nematodes and arthropods). Indeed, inferred mt-proteomes of *D. melanogaster* and *C. elegans* are about half and 65% in size, respectively, compared to those of human and mouse [14, 7, 20]. Furthermore, our recent bioinformatic analysis has suggested an even larger variation in the sizes of non-bilaterian mt-proteomes [21].

Given the large size difference in both experimentally determined and computationally predicted mt-proteomes, we asked three questions in the present study:

1. What is the mechanism behind the observed variation in mt-proteome size?
2. What is the functional significance of this variation?
3. How much of this variation can be attributed to gain/loss of Mitochondrial Targeting Signal (MTS^3^)?

The latter question concerns the predominant method of protein import into mitochondria – via the MTS/Presequence pathway. MTS refers to a targeting signal at the N-terminus end of the protein, usually within the first 90 amino-acid residues. The MTS are enriched in arginine and depleted in negatively charged residues, and form an amphipathic alpha-helical structure [22, 23] Proteins which harbor these MTS are transported into the organelle by TOM and TIM complexes in the mitochondrial outer and inner membranes, respectively. Since a large portion of the mt-proteins possess MTS, *in silico* identification of MTS is an important technique to predict protein localization.

To answer the above questions about the variation in mt-proteome composition, we analyzed the experimentally determined mt-proteomes of four bilaterian animals (*Homo sapiens* [7], *Mus musculus* [7], nematode *Caenorhabditis elegans* [13] and fruit-fly *Drosophila melanogaster* [14]) and two outgroups (*Acanthamoeba castellanii* [17] and yeast *Saccharomyces cerevisiae* [12]). We hypothesized three distinct mechanisms that could contribute to the variation in the size and content of animal mt-proteomes:

1. duplication, and/or loss of ancestral mt-proteins
2. re-localization of ancestrally non-mt-proteins to mitochondria
3. evolution of novel mt-proteins

To understand the contribution of each of these factors to the evolution of animal mt-proteomes, we subdivided all mt-proteins into four categories – ancestral proteins, proteins which underwent mitochondrial neolocalization, novel animal proteins and species-specific proteins – and analyzed the contribution of each of these categories to the mt-proteome size variation. In addition, we investigated the functional implications of this observed variation in mt-proteome size. Finally, for each of the four categories of mt-proteins, we identified the proportion of proteins possessing a MTS. Our results showed that the size-increase in mammalian mt-proteomes is primarily due to the evolution of novel animal mt-proteins in mammals and loss of ancestral mt-proteins in protostomes. In addition, some of the size difference appears to be an artifact of better mt-proteome annotation in deuterostomes. Interestingly, we found that the majority of neolocalized and novel animal mt-proteins did not possess a detectable MTS, suggesting that other import pathways play an important role in protein import of animal mt-proteins.

## 2 Methods and Materials

### 2.1 Assembling animal mt-proteomes

Four experimentally determined animal mt-proteomes were used in this study: those from *Homo sapiens, Mus musculus, Caenorhabditis elegans* and *Drosophila melanogaster*. The rat mt-proteome was not used in this analysis to avoid an over-representation of mammals. Human and mouse mt-proteomes were assembled using data from MitoCarta v2.0 [7] and IMPI (Integrated Mitochondrial Protein Index) vQ2 2018 [20]. The *C. elegans* mt-proteome was downloaded using data from Jing et al. [13]. The *D. melanogaster* mt-proteome was downloaded using data from the iGLAD database [14] (https://www.flyrnai.org/tools/glad/web/). In addition, two non-metazoan outgroups were used: *Acanthamoeba castellani* and *Saccharomyces cerevisiae*. The *A. castellani* mt-proteome was downloaded from Harvard Dataverse [17, 24] and the yeast mt-protein was downloaded from the Saccharomyces Genome Database (https://www.yeastgenome.org/) [25].

The complete proteomes for three animal species: *Homo sapiens, Mus musculus*, and *Drosophila melanogaster* and the protist *Acanthamoeba castellanii* were downloaded from Uniprot [26]. The complete yeast proteome was downloaded from the Saccharomyces Genome Database [25]. The complete proteome for *Caenorhabditis elegans* was downloaded from WormBase [27].

### 2.2 Identification of orthologous groups

Proteinortho v5.16b [28] with the default parameters (e-value:1e-05, similarity:0.25, percentage-identity:0.25, purity:1, connectivity:0.1) was used to identify groups of orthologous proteins (OGs^4^) between complete proteomes of six eukaryotic species. From the resulting sets of OGs, OGs containing mt-proteins were extracted. The OGs and the proteins contained within the OGs were subdivided into three categories, based on the presence/absence and sub-cellular localization (mitochondrial/ non-mitochondrial) of outgroup mt-proteins within individual OGs. This is described in more detail in section 3.1.1.

### 2.3 Identification of Mitochondrial Targeting Signals (MTS)

MTS were identified using MitoFates [22] (metazoan option) and TargetP v1.1 [23] (Non-Plant option). TargetP outputs a statistic (Reliability Class (RC)^5^) for each prediction, which is an indication of the strength of that prediction, with RC1 indicating the strongest and RC5 indicating the weakest prediction. MitoFates, on the other hand, provides a probability for each prediction. In this study, a protein was considered to possess a MTS if either TargetP (RC1-5) or MitoFates (any prediction probability) identified a MTS.

### 2.4 Functional analysis of mt-proteins

We used Gene Ontology (GO) analysis, pathway analysis and protein-protein interaction network analysis to assign biological function to mt-proteins. PantherDB v14.0 [29] was used for GO analysis of mt-proteins from all four animal species, while WormBase [30] and FlyMine [31] were used for analysis of *C. elegans* and *D. melanogaster* mt-proteins respectively. PantherDB was used to perform the Panther Over-representation test. For a list of given genes, the over-representation test identifies GO terms which are over-represented/under-represented when compared to a list of reference genes. In addition to GO analysis, tissue-enrichment and phenotype-enrichment analysis for genes from *C. elegans* was carried out using WormBase [32, 33].

Functional annotation of proteins is greatly aided by knowledge of their interaction-partners : protein-protein interaction networks (PPI). We used two tools, which utilize PPI information to identify biological processes and pathways enriched in different sets of mt-proteins: ConsensusPathDB Release 34 [34] and StringDB [35]. Default parameters were used for ConsensusPathDB (minimum overlap with input list:2 and p-value:0.01). ConsensusPathDB was used for functional annotation of mammalian mt-proteins, since it houses data for human, mouse and yeast. StringDB incorporates PPI data from multiple sources and identifies functional associations between query proteins. This tool was used to identify functionally-associated clusters of species-specific mt-proteins, particularly from from *C. elegans* and *D. melanogaster*.

### 2.5 Data availibility

Results of these analyses and scripts used are available on Open Science Framework (Muthye, V., Lavrov, D. (2019, July 31). Data for Causes and Consequences of animal mt-proteome size variation. https://doi.org/10.17605/OSF.IO/A49YW)

## 3 Results

### 3.1 Evolution of animal mt-proteomes

#### 3.1.1 Four categories of animal mt-proteins

We used Proteinortho v5.16b to identify groups of orthologous proteins (OGs) in four animals (*Homo sapiens, Mus musculus, Caenorhabditis elegans, Drosophila melanogaster*) and two outgroups (*Acanthamoeba castellani* and *Saccharomyces cerevisiae*). 1909 OGs had a mt-protein from at least one animal species. These were divided into three categories (Figure 1, Figure 2, Table 1):

**Table 1:** Number of OGs and mt-proteins found in each category of bilaterian animal mt-proteome. The four different categories of animal mt-proteins are outlined in Section 3.1.1.

|  | Ancestral |  | Neolocalized |  | Novel |  | Species-specific |  |
| --- | --- | --- | --- | --- | --- | --- | --- | --- |
|  | OG | mt-protein | OG | mt-protein | OG | mt-protein | OG | mt-protein |
| *C. elegans* | 322 | 344 | 192 | 233 | 233 | 238 | - | 275 |
| *D. melanogaster* | 317 | 332 | 134 | 140 | 237 | 242 | - | 124 |
| *H. sapiens* | 485 | 535 | 303 | 312 | 800 | 823 | - | 30 |
| *M. musculus* | 450 | 491 | 274 | 282 | 716 | 733 | - | 13 |

**Figure 1:**
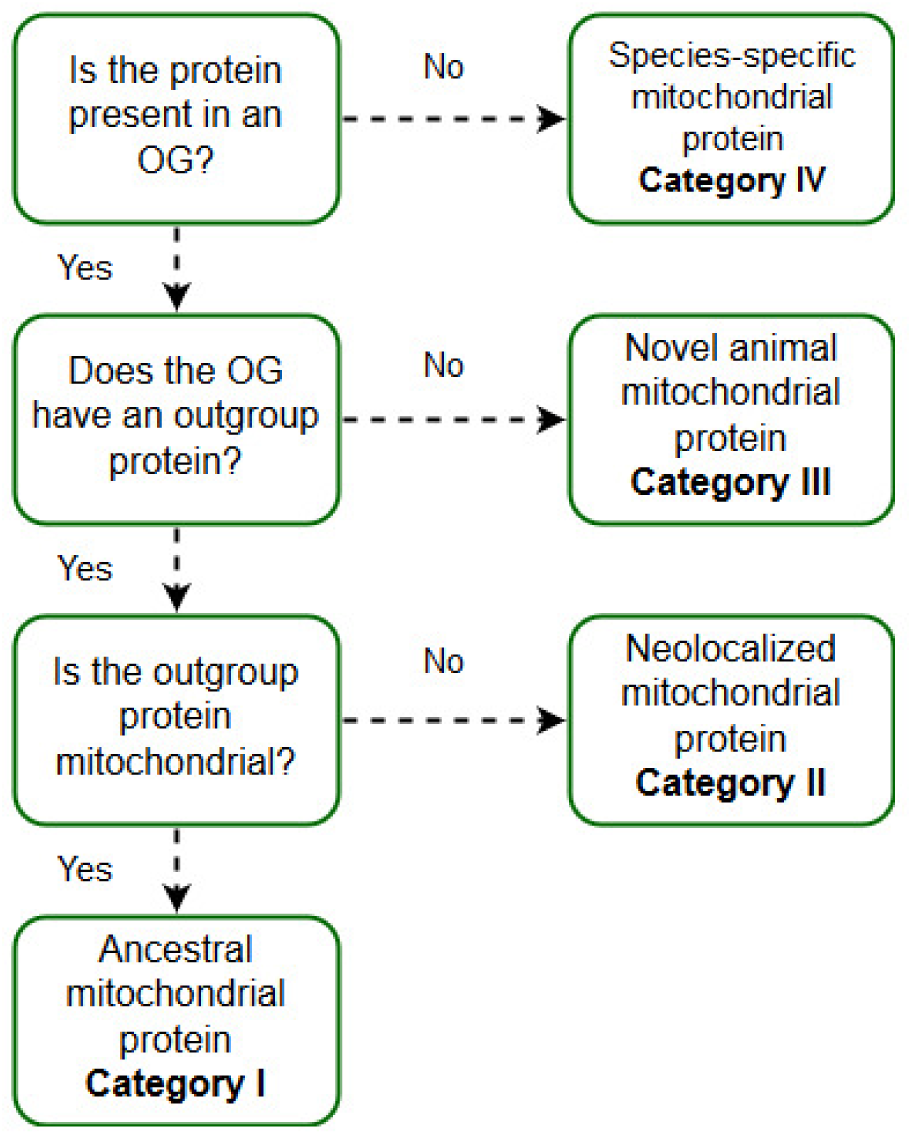
Decision tree used to subdivide animal mt-proteomes into four categories.

**Figure 2:**
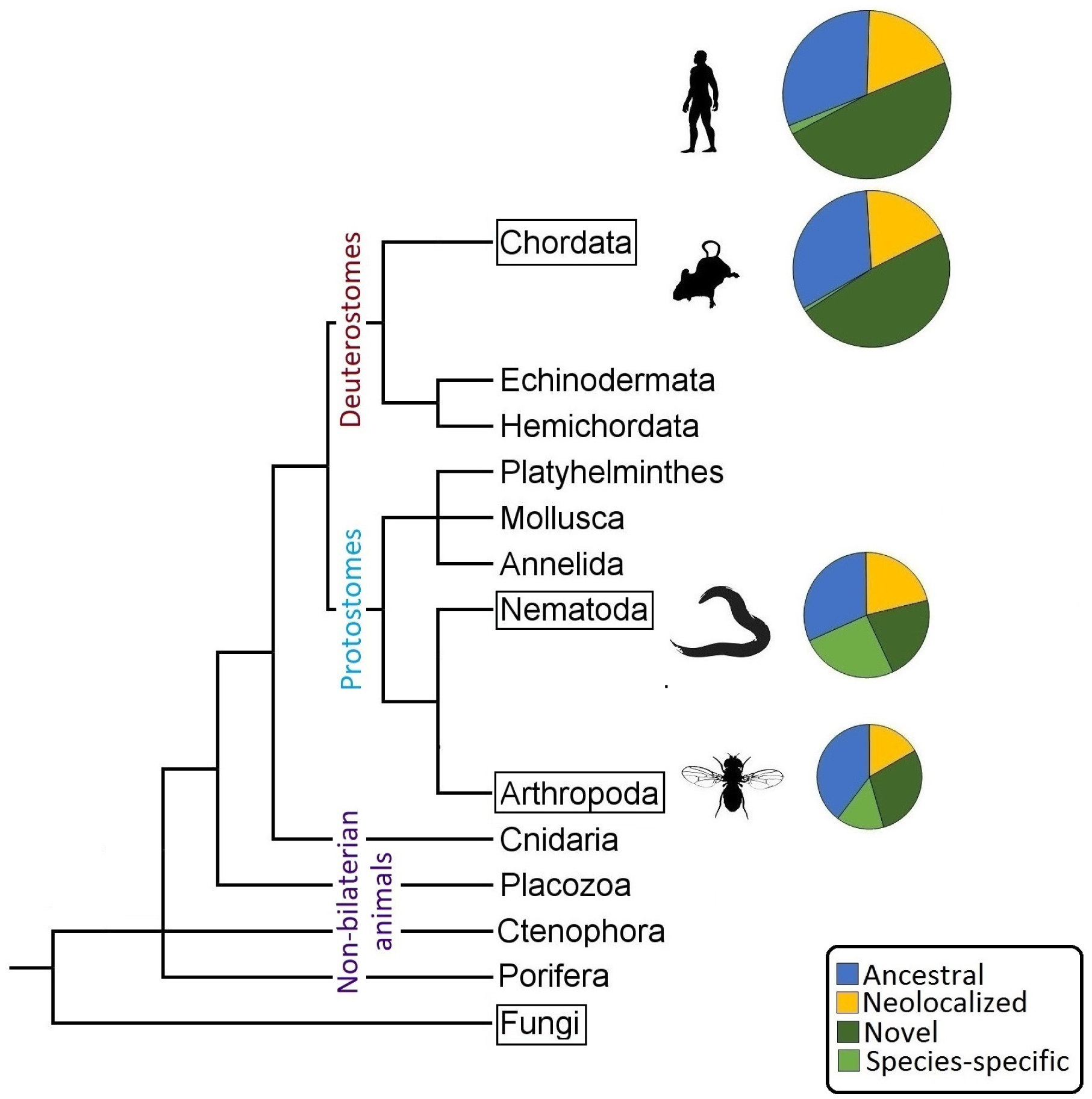
Composition of experimentally determined animal mt-proteomes. Different categories of proteins are shown by colors. The radius of the pie-chart is proportional to the overall size of the mt-proteome. The animal silhouettes were taken from PhyloPic (http://phylopic.org/). The four experimentally determined animal mt-proteomes varied in size from 838 proteins (*D. melanogaster*) to 1700 proteins (human). Animal phylogeny is modified from [36]. Taxonomic groups with characterized mt-proteomes are marked with a black box.

- Category I : OGs containing at least one mt-protein from a given animal and a mt-protein from at least one outgroup. mt-proteins within these OGs were catalogued as ancestral mt-proteins.
- Category II : OGs containing at least one mt-protein from a given animal and a non-mt-protein from at least one outgroup, but no mt-protein from either outgroups. mt-proteins within these OGs were catalogued as neolocalized mt-proteins.
- Category III : OGs containing at least one mt-protein from a given animal and no protein (mitochondrial or non-mitochondrial) from either outgroups. mt-proteins within these OGs were catalogued as novel animal mt-proteins.

Some animal mt-proteins were not a part of any OG, i.e. lacking identifiable orthologs in all other species used in the study. These were catalogued as category IV Species-specific mt-proteins. Below we describe the composition of each of the four categories of animal mt-proteins.

#### 3.1.2 Category I: Ancestral mt-proteins

In total, 528 OGs containing ancestral mt-proteins were identified. Nearly half of them (205/528) had at least one ancestral mt-protein from all four bilaterian species (Figure 3A). In addition, 239 OGs in this category were comprised of mt-proteins from both mammals but lacked a mitochondrial ortholog from either one (149) or both protostomes (90). Interestingly, the majority of OGs, lacking just one protostome mt-protein (52/74 in *C. elegans* and 60/75 in *D. melanogaster*) possessed a non-mitochondrial protein from that species, suggesting possible misannotation. Furthermore, 42 out of 90 OGs with an ancestral mt-protein from both mammals contained non-mitochondrial orthologs in both protostomes. We identified 45 and 68 proteins from *C. elegans* and *D. melanogaster*, which were present in Category I OGs and possessed a MTS, but were not annotated as mt-proteins. At the same time, 31 out of 90 OGs with an ancestral mt-protein from both mammals had no protostome protein (either mitochondrial or non-mitochondrial), indicating a loss in protostomes.

**Figure 3:**
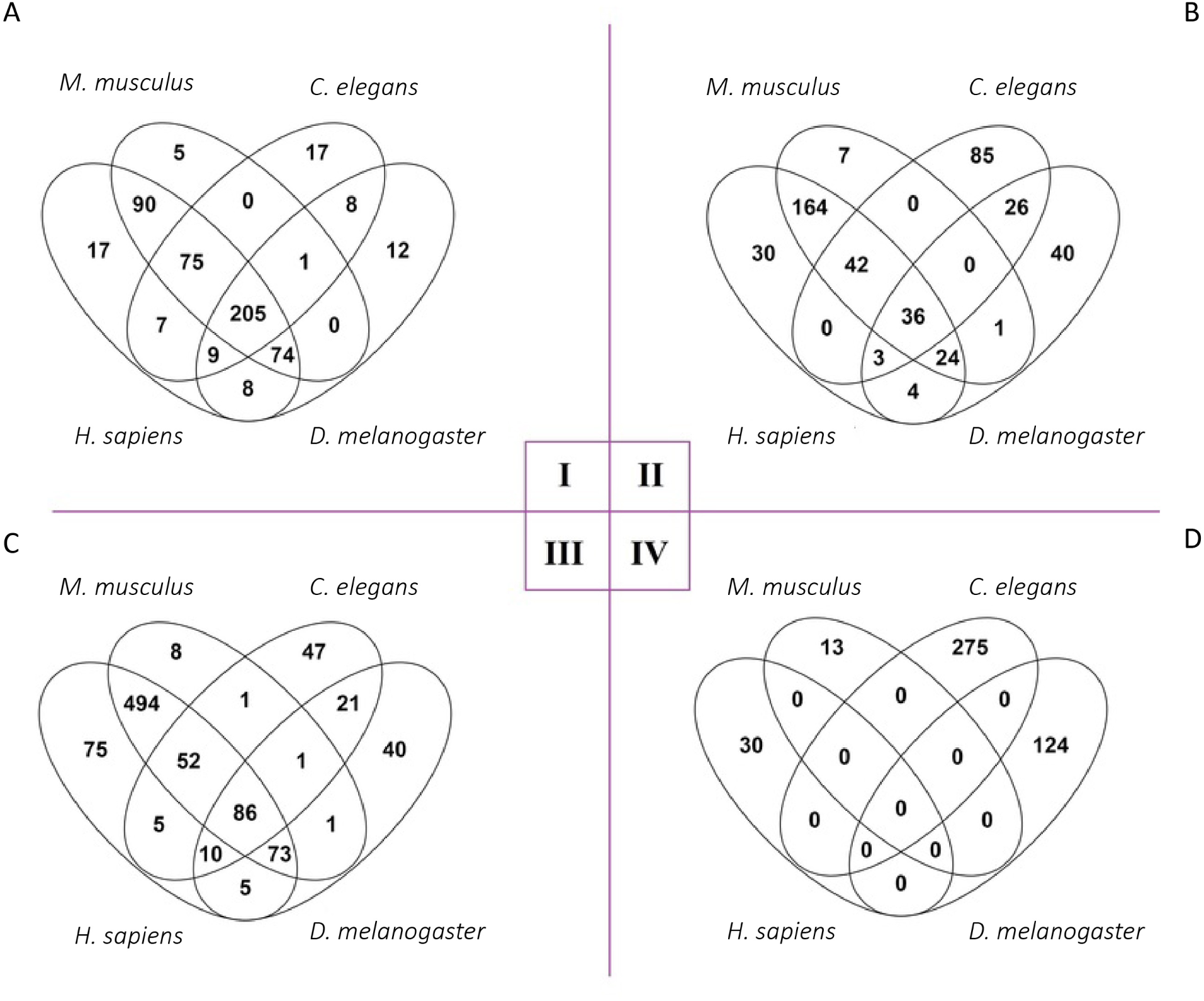
VENN diagram of Orthologous Groups (OGs) and proteins from mt-proteomes of the four animals belonging to the four categories of mt-proteins (explained in section 3.1.1) A-C] VENN diagram of OGs containing at least 1 mt-protein from categories I, II and III in the four bilaterian animals, as described in Section 3.1.1. A] Category I: ancestral mt-proteins, B] Category II: proteins which underwent mitochondrial neolocalization in animals, C] Category III: novel animal mt-proteins. D] VENN Diagram of Category IV species-specific mt-proteins from the four bilaterian animals.

#### 3.1.3 Category II: Neolocalized mt-proteins

We identified 462 OGs with proteins belonging to category II (Figure 3B). 36 OGs contained category II proteins from all four bilaterian animals, suggesting mitochondrial neolocalization prior to the protostome-deuterostome split. Additionally 66 OGs possessed mt-proteins from both mammals and one of the two protostome animals. 29/42 and 16/24 of these OGs contained an orthologous protein identified as non-mitochondrial in the other protostome species, either *D. melanogaster* or *C. elegans*, respectively. These may represent further examples of yet-to-be-identified mt-proteins. We identified 164 OGs with a category II protein from both mammals, but no mt-protein from either protostome. 123 of these 164 OGs comprised of category II proteins from both mammals and only non-mitochondrial proteins from either one or both protostomes suggesting neolocalization in mammals. The remaining 41/164 OGs contained a category II protein from both mammals and no ortholog from either of the protostomes, implying both loss and neolocalization in their evolution. Finally, in 23 category II OGs, there were mt-proteins from both protostomes, and a non-mt-protein from both mammals, suggesting a neolocalization in protostomes.

#### 3.1.4 Category III: Novel animal mt-proteins

In total, novel animal mt-proteins were identified in 919 OGs, of which only 86 (around 9%) had category III proteins from all four animals (Figure 3C). More than half of the total category III OGs (494/919) contained either a novel mitochondrial protein from both mammals but no orthologs from either of the protostomes (342 OGs) or a novel mt-protein from both mammals and a non-mitochondrial protein from either one or both protostomes (152 OGs). By contrast, only 21 OGs possessed either a novel mt-protein from both protostomes but no mammalian orthologs (10 OGs) or a novel mt-protein from both protostomes and a non-mitochondrial protein from both mammals (11 OGs).

#### 3.1.5 Category IV: Species-specific mt-proteins

13, 30, 124 and 275 category IV proteins (proteins with no identifiable orthologs) were detected in mouse, human, *D. melanogaster* and *C. elegans* respectively (Figure 3D). The large difference in the number of species-specific proteins between protostome vs. mammalian species can be explained by the difference in their estimated divergence times: ∼ 555MYA (protostomes) [37] vs. ∼ 88MYA (mammals) [38].

#### 3.1.6 Contribution of the four categories towards the size-increase of mammalian mt-proteomes

The mammalian mt-proteomes were, on average, 645 proteins larger than the protostome mt-proteomes. We quantified the extent to which each category of proteins contributed to this difference by calculating its contribution factor (CF) ^6^. For each category, the contribution factor was defined as

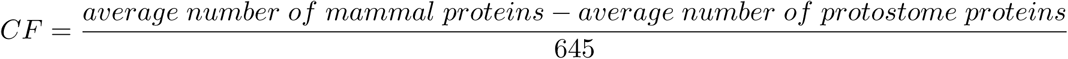

A contribution factor of 1 would indicate that the size-increase of mammalian mt-proteomes can be explained completely by that category, while 0 would indicate that category has no effect. A negative value would indicate a higher number of protostome proteins in that category compared to mammals. Since category IV is a special case of category III, we combined the contribution factors for both categories into a single score. The contribution factors were 0.17, 0.27 and 0.56 for category II, I and (III+IV) respectively.

In addition to neolocalization and evolution of novel mt-proteins, we also hypothesize that duplication of genes encoding mt-proteins could also result in an increase in size of mammalian mt-proteomes. To estimate the contribution of gene-duplication towards mitochondrial proteome size-variation, we calculated the contribution factor per category as follows:

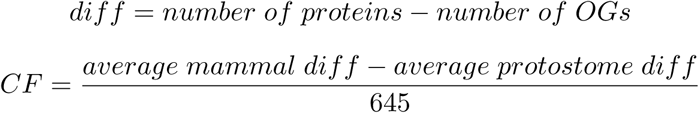

The contribution factors for gene-duplication in the three categories I-III were low (0.04 (category I),-0.02 (category II) and 0.02 (category III) respectively).

### 3.2 Functional analysis of mt-proteins

### 3.2.1 Expansion of pathways in mammalian mitochondria

On average, mammals had 538 more novel animal mt-proteins (category III) compared to protostomes. In mammals, majority of these novel animal mt-proteins were present in both human and mouse (494 OGs). Several proteins from these 494 OGs were additions to the mitochondrial inner (44) and outer (54) membranes, locations of critical mitochondrial functions such as energy metabolism and metabolite and protein import. Panther Over-representation test showed that these mammal-specific novel animal mt-proteins were involved in mitochondrial translation, apoptosis, complex I assembly and oxidative phosphorylation. “Thermogenesis”, “Butanoate metabolism”, “Apoptosis” and “Oxidative phosphorylation” were the top enriched pathways detected by ConsensusPathDB.

Novel mammalian mt-proteins could represent either novel biological pathways or additions to existing pathways. KEGG analysis revealed that 86% (231/268) of pathways identified in at least one animal species were present in all four animals. Thus, most of the mammal-specific mt-proteins were additions to existing animal mitochondrial pathways. In nearly half (118/268) of the identified pathways, the average number of mammalian proteins was at least double the number of invertebrate proteins. Majority of them were signaling pathways, like “JAK-STAT signaling pathway”, “Prolactin signaling pathway” and “ErbB signaling pathway”.

#### 3.2.2 Loss of proteins involved in amino-acid metabolism in protostomes

528 OGs included ancestral mt-proteins, of which 31 OGs contained ancestral mt-proteins from both mammals, and no orthologs in either protostome, thus representing loss in protostomes. 24 proteins from these 31 OGs were mapped to at least 1 pathway by ConsensuspathDB including metabolism of amino acids and derivatives (8) and urea cycle (3). Other ancestral proteins lost in protostomes included Monocarboxylate transporter 1 (SLC16A1), Magnesium transporter (MRS2) and Protein adenylyltransferase SelO (SELENOO).

#### 3.2.3 Protostome-specific mt-proteins

We identified 507 protostome-specific mt-proteins : novel animal proteins (category III) and species-specific proteins (category IV) in the two analyzed protostome mt-proteomes. Only 21 of them were present in both species. 322 *C. elegans* proteins did not have any mitochondrial orthologs in other species and 275 of them had neither mitochondrial nor nuclear orthologs (category IV). Similarly, 164 *D. melanogaster* proteins did not have mitochondrial orthologs in other species, of which 124 lacked both mitochondrial and nuclear orthologs (category IV).

Interestingly, many of these 275 category IV proteins from *C. elegans* were involved in nematode-specific functions, like nematode larval development, embryo development and protection against microbes. Phenotype-enrichment analysis identified “innate immune response variant” and “avoids bacterial lawn” as the top enriched terms. StringDB identified 55 clusters of mt-proteins with at least 2 interacting members. The largest cluster contained 16 proteins involved in “lipid metabolism” and “nutrient reservoir activity”. The next largest cluster of 9 proteins were possible sperm mt-proteins. The remaining three clusters comprised of DNA-binding proteins, proteins involved in regulation of transcription, mitochondrial membrane proteins and proteins involved in ATP hydrolysis coupled proton transport.

In *D. melanogaster*, 124 mt-proteins did not have an ortholog in any of the other species in this study. As per PantherDB, the top enriched protein classes of these *D. melanogaster*-specific mt-proteins were “oxidoreductases” (21) and “proteases” (14). FlyMine identified pathways for 41/124 *D. melanogaster* proteins. These proteins were involved primarily in metabolic pathways, with 11 involved in oxidative phosphorylation and 9 in lipid metabolism. StringDB identified 14 clusters of mt-proteins with at least 2 interacting members. The largest cluster possessed 10 proteins involved in sperm mitochondrial processes. These include Sperm-Leucylaminopeptidase 3, Sperm-Leucylaminopeptidase 5, Sperm-Leucylaminopeptidase 8 and Loopin-1.

### 3.3 Role of MTS in mt-proteome evolution

The proportion of mt-proteins with a MTS ranged from 41% to 56% among the four animal species. However, because of the difference in the mitochondrial proteome size, the absolute number of proteins with MTS was nearly twice as large in *H. sapiens* compared to *C. elegans* and *D. melanogaster*. We noticed a significant difference in the proportion of proteins possessing a MTS among the four categories of proteins described above (Figure 4). The average proportion of proteins with MTS ranged from 30% in category II to 66% in category I. For category I, III and IV, the proportion of proteins with MTS was lowest in *C. elegans* and highest in *D. melanogaster* (Table 2).

**Table 2:** Proportion of proteins possessing a MTS from individual categories of the bilaterian animal mt-proteomes

|  | Category I | Category II | Category III | Category IV |
| --- | --- | --- | --- | --- |
| *C. elegans* | 63.95% | 15.88% | 47.06% | 29.09% |
| *D. melanogaster* | 68.07% | 28.57% | 54.55% | 57.26% |
| *H. sapiens* | 66.73% | 40.06% | 48.48% | 40.00% |
| *M. musculus* | 67.01% | 37.59% | 48.16% | 38.46% |

**Figure 4:**
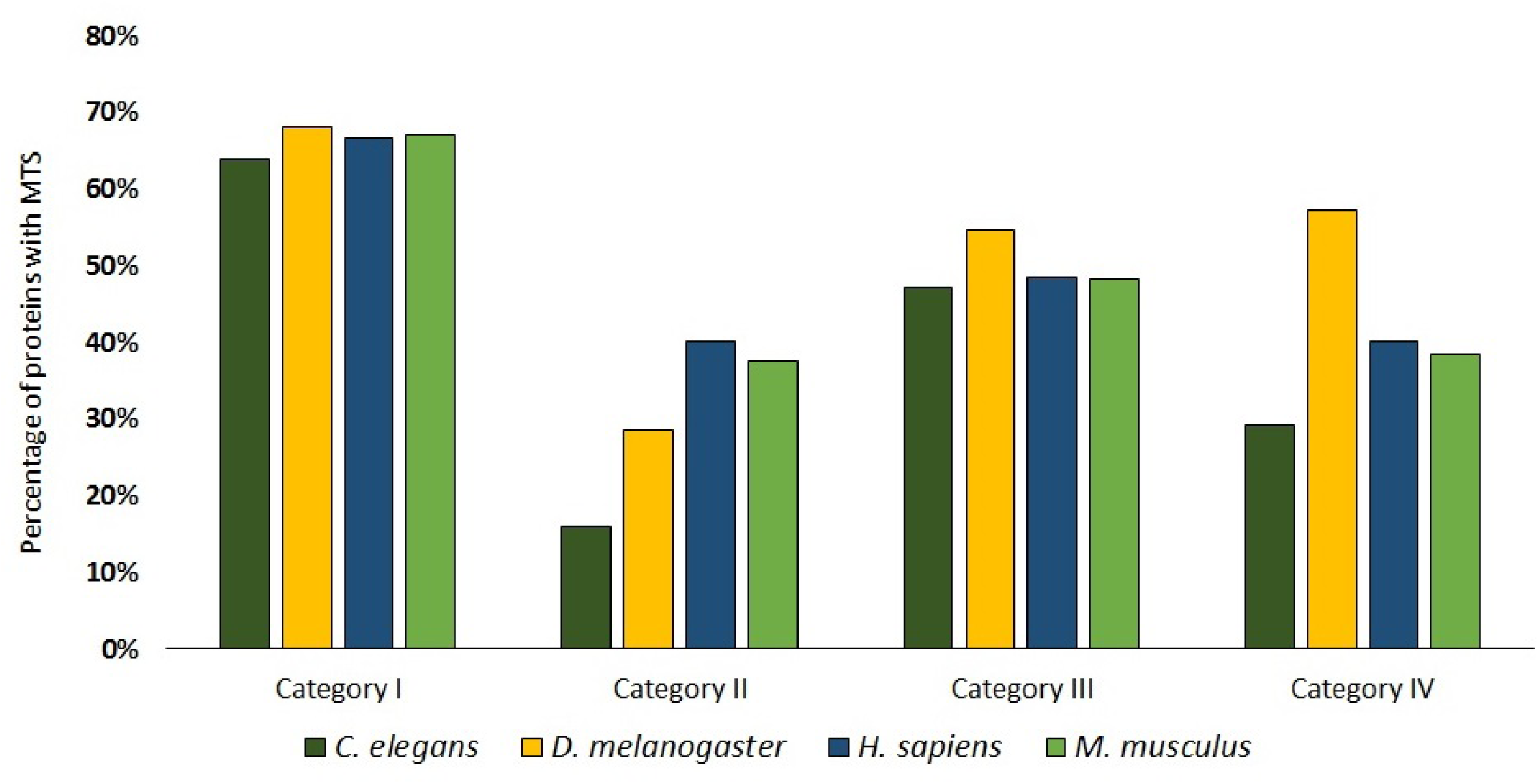
Percentage of proteins with MTS in all the four categories in animal mt-proteomes.

## 4 Discussion

One might expect that the centrality of mitochondria in multiple crucial cellular processes in animals would result in a large core of conserved animal mt-proteins with few lineage-specific additions/deletions. This expectation is reinforced by the near-identical mtDNA gene-content of the four animals used in this study [39, 40, 41, 42]. By contrast, our comparative analysis of the four animal mt-proteomes paints a more dynamic picture of the organellar proteome – a small core of ancestral conserved mt-proteins and a large number of lineage-specific differences (mammal-specific, nematode-specific or arthropod-specific) (Figure 2).

On average, while just 19% of each animal mt-proteome was conserved in all four animals, 42% did not have mitochondrial orthologs in other species in this study, i.e. lineage-specific mt-proteins. There are two possible interpretations for these results: 1] observed differences in mt-proteome reflect morphological and ecological differences among the animal species used for the study, or 2] observed differences can be explained by our inability to detect orthologous proteins in faster evolving protostome genomes. While both of these interpretations can be valid to some extent, functional analysis of observed differences favour the former one. For example, a large number of mammal-specific mt-proteins were involved in signaling pathways, a finding consistent with increased morphological and cellular complexity of mammals. The importance of mitochondria as signaling organelle in mammals is also well-supported by recent studies [43, 44].

By contrast, in the nematode, interactions with its surrounding microbiome have influenced some of changes in the mt-proteome, with several species-specific mt-proteins involved in protection of the nematode from bacteria and fungi. *C. elegans* is a free-living nematode which grows on decomposing materials containing a high number of micro-organisms [45]. These microbes play a critical role in the life of the nematode, as food, competitors or even as pathogens and parasites. Interestingly, mitochondria themselves may be targeted by pathogenic microbes, and several proteins have been identified which up-regulate protective pathways following such mitochondrial disruption [46].

In addition to lineage-specific gains, another factor contributing to the variation in size and content of animal mt-proteomes was the loss of ancestral mt-proteins in protostomes. Indeed, ecdysozoans are known to have undergone extensive gene-loss [47]. For instance, in both protostomes, enzymes involved in amino-acid metabolism, such as those involved in arginine biosynthesis were lost, suggesting that these enzymes were dispensable in *C. elegans* and *D. melanogaster*. These enzymes have been lost independently in multiple eukaryotic lineages [48]. The dual-role of these enzymes in the urea-cycle in vertebrates seems to have been responsible for the conservation of the arginine biosynthesis pathway in vertebrates.

Our analysis showed that most of the novel genes encoding mt-proteins are lineage-specific (mammal-specific, *C. elegans*-specific or *D. melanogaster*-specific), whereas comparatively few genes evolved before the protostome-deuterostome split. This result contrasts previous studies on the comparative analysis of animal genomes, which showed that most of the genomic novelty originated in the last common metazoan ancestor, and at the origin of bilaterian animals with few lineage-specific changes [49, 50]. This discrepancy can be due to the limited taxonomic sampling of our study. If some novel mt-proteins were acquired either in the common ancestor of all metazoans or in the lineage leading to bilaterian animals and then lost in Ecdysozoa, the pattern of presence/absence of such proteins would be indistinguishable from the gain of these proteins in deuterostomes. The other major protostome group, lophotrochozoans, did not experience a gene-loss as severe as the ecdysozoans [51]. Characterization of mt-proteomes from this group can help differentiate whether a mt-protein was gained in mammals or lost in protostomes. In deuterostomes, both mt-proteomes belong to a single class (Mammalia) from a single phylum (Chordata). Also, experimental characterization of animal mt-proteomes is limited to bilaterian animals, which represent just one of the five major lineages of animal phylogeny. Thus, experimental characterization of mt-proteomes from more animal lineages, such as lophotrochozoans and non-bilaterian animals, would aid in analysis of the metazoan mt-proteome. Additionally, it is also possible that some of the variation among mt-proteomes is due to the technical problems with orthology detection. While generating OGs, Proteinortho might miss some of the faster-evolving orthologs in *C. elegans* or *D. melanogaster*. This might result in overestimation of the number of lineage-specific mt-proteins.

Nearly, 1/5th of the animal mt-proteins are mitochondrial neolocalized proteins. Interestingly, majority of the mitochondrial recruitments did not happen via gaining a MTS. MTS play an important role in import of ancestral mt-proteins, but majority of mt-proteins from other classes lacked MTS. This suggests that 1] alternate targeting pathways or non-canonical MTS facilitate import of a sizeable portion of animal mt-proteins and 2] an MTS-only approach to identify novel mt-proteins would miss a large number of proteins in this category. To date, several alternate pathways which import proteins in the mitochondria have been identified. For example, all of the mitochondrial outer-membrane proteins and majority of the intermembrane-space proteins contain non-canonical signals within the mature protein [4]. In some proteins, like DNA helicase Hmi1p in yeast, the targeting signal is present at the C-terminal end of the protein instead of the canonical N-terminal end [52]. Identification of such alternate targeting signals and development of tools to identify these signals in proteins can further our understanding of mt-protein import.

Some additional factors need to be considered while interpreting results of this study, like differences in methods used in characterization of mt-proteomes and variation in completeness of individual animal mt-proteomes. The four animal mt-proteomes were characterized using different experimental and bioinformatic approaches. Each of these methods have their pros and cons. For instance, MS based methods face the problem of co-purifying contaminants and missing low-abundance mt-proteins. On the other hand, bioinformatic methods like detection of MTS, would miss non-canonical targeting signals. Mammalian mt-proteomes represent the most complete mt-proteomes of the four animals, because both experimental and computational techniques were applied simultaneously to identify mt-proteins. However, even the best studied mt-proteomes (human and mouse) are suggested to be only around 85% complete [6].

## 5 Conclusion

Animal mt-proteomes vary with respect to size and content. Using computational techniques, we investigated the causes and functional significance of this variation in size and content of animal mt-proteomes. We found that several factors contributed to the above-mentioned size difference: evolution of novel mt-proteins, loss of ancestral mt-proteins, neolocalization of non-mt-proteins to mitochondria and duplication of genes encoding mt-proteins. Evolution of novel mt-proteins was the primary reason for the increase in size of mammalian mt-proteomes. Functional analysis of these novel mammalian mt-proteins suggest an increased role of mitochondria in cellular signaling. In addition to novel biological functions, these novel mammalian mt-proteins were also additions to existing mitochondrial pathways and complexes. Loss of ancestral mt-proteins in protostomes, like those involved in amino-acid metabolism, also resulted in the size-variation in animal mt-proteomes. Surprisingly, gain of MTS was not a significant contributor to the size-variation. In fact, majority of novel and neolocalized mt-proteins in animals lack an identifiable MTS, suggesting either an increased role of alternate mt-protein import pathways or presence of non-canonical targeting signals which are not identified by existing methods. Experimental characterization of mt-proteomes from other animal lineages, from both bilaterian and non-bilaterian phyla, and development of tools to identify non-canonical mitochondrial targeting signals will help further our understanding of animal mt-proteome evolution.

## 6 Acknowledgement

We gratefully acknowledge Dr. Carolyn Lawrence-Dill, Dr. Karin Dorman, Dr. Iddo Friedberg and Dr. Robert Jernigan for their comments and suggestions regarding the manuscript.

## Footnotes

1 mt-proteins : mitochondrial proteins

2 mt-proteomes : mitochondrial proteomes

3 MTS : mitochondrial targeting signal

4 OG: Orthologous Group

5 RC: Reliability Class

6 CF: Contribution Factor

